# Tdrd6a regulates the aggregation of Buc into functional subcellular compartments that drive germ cell specification

**DOI:** 10.1101/267971

**Authors:** Elke F. Roovers, Lucas J.T. Kaaij, Stefan Redl, Alfred W. Bronkhorst, Kay Wiebrands, António M. de Jesus Domingues, Hsin-Yi Huang, Chung-Ting Han, Willi Salvenmoser, Dominic Grün, Falk Butter, Alexander van Oudenaarden, René F. Ketting

## Abstract

In recent years, it has become clear that phase separation represents an important class of subcellular compartmentalization. However, relatively little is known about how the formation or disassembly of such compartments is regulated. In zebrafish, the Balbiani body (Bb) and the germ plasm (Gp) are phase-separated structures essential for germ cell specification and home to many germ cell-specific mRNAs and proteins. Throughout development, these structures range from a single large aggregate (Bb), to a dispersed state and back to relatively large assemblies (Gp). Formation of the Bb requires Bucky ball (Buc), a protein with prion-like properties. We found that the multi-tudor domain-containing protein Tdrd6a interacts directly with Buc, affecting its mobility and aggregation properties. Importantly, lack of this regulatory interaction leads to significant defects in germ cell development. Our work presents a new mechanism for how prion-like protein-aggregations can be regulated and highlights the biological relevance of such regulatory events.

## Introduction

Phase-separating mechanisms are increasingly being acknowledged as important aspects of cell biology. After the initial description of the liquid-like behavior of P-granules, peri-nuclear RNA-rich protein aggregates in the *C. elegans* germline, many other RNA-containing ‘granules’ have been shown to have similar properties (Brangwynne et al., 2009, 2011; Kroschwald et al., 2015). Important players in the formation of these structures are proteins containing intrinsically disordered regions (IRDs) and/or prion-like domains (PrDs) (Kato et al., 2012; Kroschwald et al., 2015). Such proteins have the propensity to self-aggregate and potentially take along other proteins in the process (Prusiner, 1998; Shorter and Lindquist, 2005). In many ways, biologically functional protein assemblies such as P-granules resemble pathogenic protein-aggregation states. It has been suggested that such disease-causing aggregations are an extreme manifestation of an abundantly used mechanism to form membrane-less compartments (Shin and Brangwynne, 2017). This suggests that mechanisms are in place that prevent healthy, functional aggregates to transform into pathological forms. However, insights into such regulatory mechanisms have remained scarce.

In many organisms germ cell fate is imposed on embryonic cells through the cytoplasmic inheritance of P-granule-like structures, called germ plasm (Gp) (Ikenishi, 1998; Raz, 2003). In zebrafish, Gp originates from an evolutionary conserved electron-dense aggregate in the oocyte, called the Balbiani body (Bb) (Kloc et al., 2004). The mRNAs enriched in the Bb and Gp are often germline-specific and in zebrafish these include *vasa*, *nanos3* and *dazl* (Hashimoto et al., 2004; Koprunner et al., 2001; Yoon et al., 1997). Depletion of single Gp mRNAs can have detrimental effects on primordial germ cell (PGC) numbers, showing that individual Gp components are important for PGC specification and survival (Koprunner et al., 2001; Slaidina and Lehmann, 2017; Tzung et al., 2015; Weidinger et al., 2003).

Thus far, Bucky ball (Buc) is the only protein that is known to play a key role in the formation of the Bb in zebrafish (Bontems et al., 2009; Marlow and Mullins, 2008). Overexpression of Buc in zygotes revealed that Buc is sufficient to induce ectopic PGCs, suggesting it is also involved in the formation of the Bb-related Gp structure (Bontems et al., 2009). Buc is a protein with a PrD and elegant studies on its homolog in *Xenopus* (Xvelo) have demonstrated that these proteins self-aggregate into membrane-less organelles that display amyloid-like features (Boke et al., 2016).

Core piRNA-pathway components, such as Ziwi in zebrafish and Aub in *Drosophila*, are present in the Gp as well (Harris and Macdonald, 2001; Houwing, 2009). Furthermore, it has been shown in *Drosophila* that piRNA-pathway components inherited via the Gp are essential for transposon silencing in the offspring (Brennecke et al., 2008), and piRNA-mRNA interactions have been proposed to drive mRNA localization to Gp (Barckmann et al., 2015; Vourekas et al., 2016). Many proteins involved in the piRNA-pathway have been identified through genetic and biochemical approaches including multi-Tudor domain-containing proteins (Tdrds) (Siomi et al., 2010). Tdrds play important roles in the formation of nuage, a peri-nuclear protein-RNA aggregate that associates closely with mitochondria. For some Tdrds it has been shown that they bind to symmetrically dimethylated arginine (sDMA) residues on their interaction partners. In zebrafish for instance, the interaction between Tdrd1 and the Piwi protein Zili in zebrafish is mediated via a specific sDMA site in Zili (Huang et al., 2011).

One of the Tdrds that has received relatively little attention, is Tdrd6, the closest vertebrate homolog to *Drosophila* Tudor (Tud). Tud has been shown to interact with Piwi proteins Aub and Ago3 and plays a role in localization of Aub to Gp and polar granule formation (Kirino et al., 2010; Nishida et al., 2009; Thomson and Lasko, 2004). In mice, TDRD6 plays a role in establishing the chromatoid body, a testis-specific structure that resembles Gp, and the localization of piRNA pathway components to this body (Vasileva et al., 2009). In addition, it has been reported to be involved in spliceosome assembly in primary spermatocytes (Akpınar et al., 2017). However, a specific molecular function of Tdrd6 or Tud has thus far not been demonstrated.

In this study, we show that that Tdrd6a is required for coordinated loading of essential components into PGCs through fine-tuning of the aggregating properties and mobility of the Bb organizer Buc. As such, the Tdrd6a-Buc interaction represents one of the few documented cases of how the aggregation of a prion-like protein can be regulated *in vivo*, such that it can fulfil an important biological function without causing detrimental aggregation. Considering the presence of Tudor domains in various proteins, we speculate that similar mechanisms act in other cell types that have been related to protein-aggregation-linked diseases.

## Results

### Tdrd6a is gonad-specific and localizes to nuage, the Bb and Gp

The zebrafish genome encodes three Tdrd6 paralogs: *tdrd6a-c*. In this study, we focused on *tdrd6a*. Tdrd6a contains seven Tudor domains and is 2117 amino acids in length (Figure S1A). Germline-specific expression of *tdrd6a* was validated by RT-PCR (Figure S1B). Immunohistochemistry (IHC) confirmed that Tdrd6a is expressed in the ovary, where Tdrd6a localizes to nuage (Figure 1A, arrowhead) and to the Bb (Figure 1A, arrow). Tdrd6a is also maternally provided and localizes to the Gp in 4-cell stage embryos (Figure 1B, arrowheads). 24 hours post fertilization (hpf), Tdrd6a is restricted to PGCs, where it again localizes to nuage (Figure 1C, arrowheads). We confirmed the identity of the Tdrd6a-containing structures using established markers for the nuage, Bb and the Gp (Figure S1C). In summary, these results demonstrate that Tdrd6a is maternally contributed and localizes to three conserved and related structures involved in germline specification and maintenance: the Bb, Gp and nuage.

**Figure 1.**
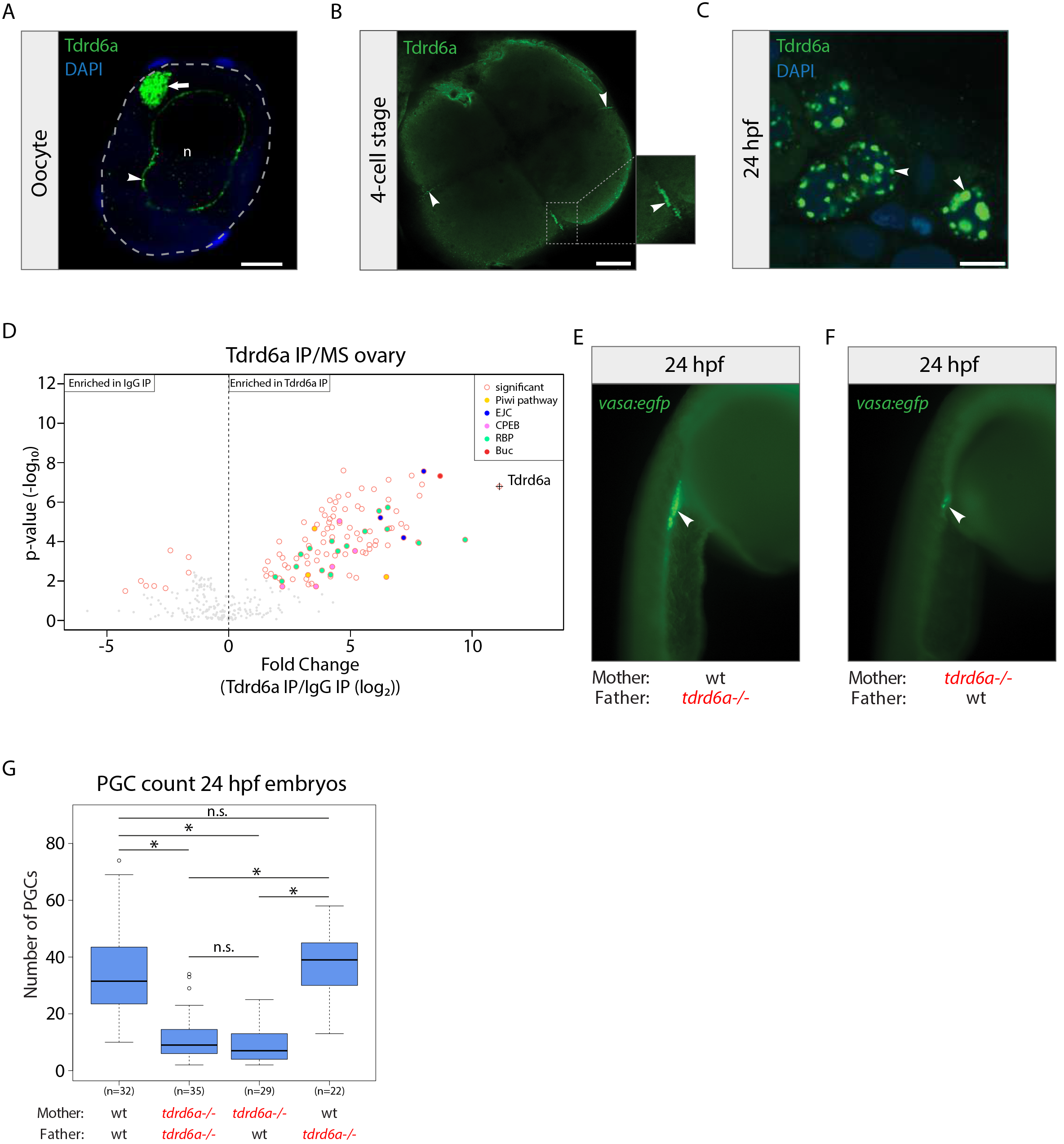
Tdrd6a is germline specific and required for PGC formation. (A) IHC for Tdrd6a (green) in oocytes. Blue: DAPI. Arrowhead and arrow indicate Tdrd6a staining in the nuage and Bb, respectively. Grey dashed line outlines the cell and n indicates the nucleus. Scale bar = 10μm. (B) IHC for Tdrd6a (green) in 4-cell stage embryos. Arrows indicate Tdrd6a localization to the Gp. Scale bar = 100μm. (C) Tdrd6a (green) localizes to perinuclear nuage granules 24hpf (arrowheads). Blue: DAPI. Scale bar: 7.5 μm. (D) MS of Tdrd6a IPs on ovary, compared to IgG control. (E,F) 24hpf embryos derived from a wt (E) or mutant mother (F) in a *vasa:egfp* background. Arrowheads indicate the PGCs. (G) Quantification of PGC numbers in 24hpf embryos from the crosses indicated on the X-axis (* indicates p-value < 0.0001, n.s. indicates non-significant, calculated by Wilcoxon test).

### Identification and characterization of a *tdrd6a* mutant allele

We isolated a *tdrd6a* allele harboring a premature stop codon (Q158X) from an ENU mutagenized library as described (Wienholds, 2002). Western blot analysis confirmed loss of Tdrd6a in homozygous mutant animals (Figure S1D). Homozygous zygotic (Z) and maternal-zygotic (MZ) *tdrd6a* mutants are fertile, indicating that Tdrd6a is not essential for fertility. *Tdrd6a−/−* oocytes show complete loss of Tdrd6a staining in perinuclear nuage (Figure S1E, arrowhead) and Gp in 4-cell stage embryos (Figure S1F, arrowheads). Some residual staining remains in the Bb in *tdrd6a* mutants (Figure S1E, arrow), however, a strong Tdrd6a-related Bb phenotype (see later) suggests that this is due to cross reactivity of the antibody. We conclude that *tdrd6a*^*Q158X*^ represents a strong loss of function allele.

### Tdrd6a interacts with Ziwi but does not affect piRNA populations

Next, we performed a Tdrd6a immunoprecipitation (IP) on ovary lysates, followed by label-free quantitative mass spectrometry (LFQ-MS). Besides Tdrd6a we found strong enrichments for several complexes containing RNA-binding proteins (RBPs), including the Exon Junction Complex (EJC) and the CPEB complex (Figure 1D). In addition, we identified the Piwi pathway components Ziwi, Zili, and Tdrd7 (Figure 1D). Finally, we found that Bb component Buc was highly enriched (Figure 1D).

Given the interaction with Ziwi and Zili, we probed for a role of Tdrd6a in the piRNA-pathway. We first performed Tdrd6a co-IP experiments followed by Western blot analysis for Zili and Ziwi. In comparison to Tdrd1 (Huang et al., 2011), Tdrd6a primarily interacts with Ziwi (Figure S1G). Next, we studied the impact of Tdrd6a on piRNAs. Small RNA (smRNA) sequencing of total ovary did not show significant differences between *tdrd6a+/−* and *tdrd6a−/−* animals. General piRNA characteristics, including length distribution, ping-pong signature, 5’U bias and transposon-targeting repertoire were unaffected. However, we did observe a small but significant reduction in the typical antisense bias for piRNAs mapping to retro-transposons (Figure S1H-K). When we roughly divided oocytes into early (ø < 300 μm) and later stage oocytes (ø > 300 μm), we noticed that this represents a defect in accumulation of antisense piRNAs during early oogenesis only (Figure S1L-S1P). In conclusion, while Tdrd6a associates with Ziwi, its absence has little effect on piRNA activity in zebrafish.

### Tdrd6a affects PGC formation

MZ *tdrd6a* mutants have a strong tendency to develop into males. Since the amount of primordial germ cells (PGCs) can have an impact on sex-determination in zebrafish (Tzung et al., 2015), we examined the effect of Tdrd6a on PGC formation. In both wt and MZ *tdrd6a−/−* embryos, PGCs marked by the *vasa:egfp* transgene (Krøvel and Olsen, 2002) are at the genital ridge at 24hpf (Figure 1E and F, arrowhead). However, we observe a significant reduction in PGC number in the offspring from *tdrd6a−/−* females, irrespective of the genotype of the father (Figure 1G). We did not observe ectopic PGCs, indicating that Tdrd6a does not significantly affect PGC migration.

### ScRNA-seq reveals a developmental delay in MZ *tdrd6a−/−* PGCs

To learn more about the underlying cause of the PGC defect, we performed single-cell RNA-sequencing (scRNA-seq) on PGCs isolated from embryos spawned by *tdrd6a+/−* (hereafter called: wt) and *tdrd6a−/−* (hereafter called: Mmut) mothers crossed with wt males. PGCs were marked using the *kop:egfp-f-nos1-3’UTR* transgene (Blaser et al., 2005) and isolated by fluorescence-activated cell sorting (FACS) (Figure 2A). Three time-points were analyzed: 1) when PGCs can be first identified using transgenic GFP expression (3.5hpf), 2) during migration of the PGCs (8hpf) and 3) when the PGCs have reached the genital ridge (24hpf).

**Figure 2.**
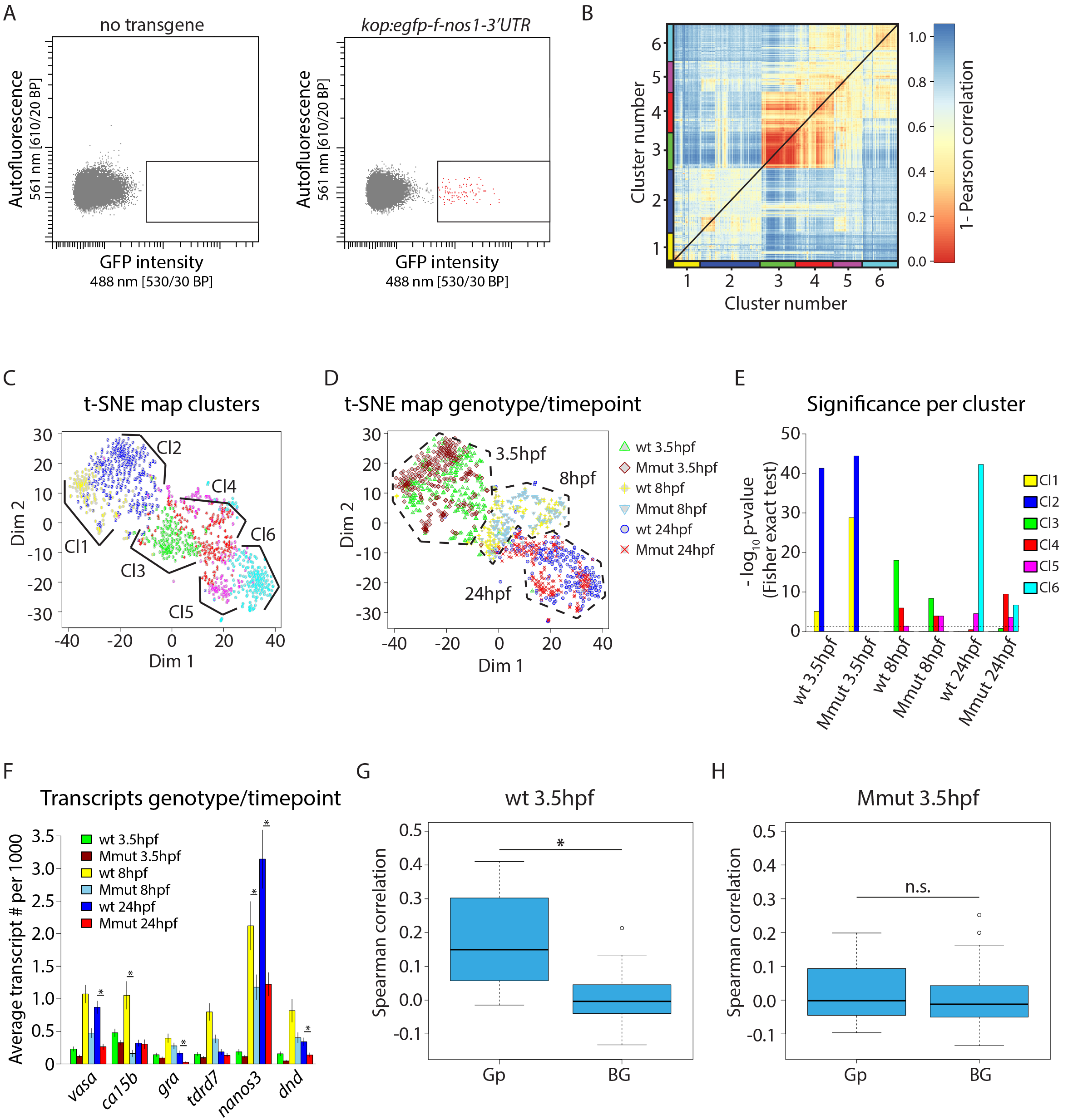
Single cell RNA-seq analysis reveals that maternal Tdrd6a mediates positive correlation of Gp residing mRNAs loading into PGCs. (A) Flow cytometry plots of the sort strategy used in this study. Representative FACS plots of embryos 8hpf without or with the *kop:egfp* transgene are shown. Positive events are indicated in red. (B) Heatmap indicating transcriptome distances of ~1100 PGCs computed as 1-Pearson’s correlation coefficient. K-medoids clustering identified six clusters, which are color coded on the x and y-axis. (C) Similar as in (B), but now visualized in a t-SNE map. Clusters identified by k-medoids clustering are color coded as in (B). (D) t-SNE map highlighting the genotype and developmental timepoint of the individual PGCs as indicated. (E) Barplot displaying the significance of enrichment for the different genotype-developmental time combinations in the six clusters identified in (B). (F) Barplot showing the average transcript counts per 1000 transcripts per cell of six Gp transcripts in all six different genotype-developmental timepoint combinations, as indicated. Error bars represent the SEM (* indicates p-value < 0.01, calculated by negative binominal statistics and corrected for multiple testing (Benjamini-Hochberg)). (G, H) Boxplots displaying the Gp-Gp and BG-BG correlations in wt and Mmut embryos, respectively (* indicates p-value < 0.001, n.s. indicates non-significant, calculated by Wilcoxon test).

We used a modified version of the original CEL-seq protocol to sequence ~1100 individual PGCs (Grün et al., 2014; Hashimshony et al., 2012) (Figure S2A) and analyzed the data using RaceID2 (Grün et al., 2016). Representation of the pairwise distances of the single cell transcriptomes in a heatmap revealed two main clusters, which can be further subdivided into clusters 1-2 and clusters 3-6 by k-medoids clustering (Figure 2B). Representation of this data in t-distributed stochastic neighbor embedding (t-SNE) maps (Van Der Maaten and Hinton, 2008), color-coding every cell based on the cluster they belong to (Figure 2C) and indicating the genotypes of the individual cells and their developmental timepoints (Figure 2D), revealed that clusters 1-2 predominantly harbor 3.5hpf old PGCs, whereas clusters 3-6 consist of PGCs from 8hpf and 24hpf (both wt and Mmut). Consistent with this, the pluripotency gene *nanog* is selectively expressed in clusters 1-2, comprised mostly of 3.5 hpf cells (Figure S2B) (Takahashi and Yamanaka, 2006). In contrast, the *rps* gene family, which has been shown to be upregulated after the MZT (Siddiqui et al., 2012), is upregulated in cluster 3-6, containing mostly 8hpf and 24hpf cells (Figure S2C).

Next, we computed the enrichment and the fraction of cells of the different genotype-developmental time-point combinations in the identified clusters. For cells sorted at 3.5hpf and 8hpf no strong differences between genotypes could be observed (Figure 2E and Figure S2D). However, a significant fraction of Mmut 24hpf PGCs was enriched in cluster 4 (Figure 2E), which is dominated by 8hpf old PGCs. This suggests that PGCs lacking maternal Tdrd6a become developmentally delayed between 8hpf and 24hpf.

### Tdrd6a affects coordinated loading of Gp mRNAs into PGCs

Since individual Gp transcripts can influence PGC numbers (Koprunner et al., 2001; Tzung et al., 2015; Weidinger et al., 2003), we tested if Mmut PGCs generally have lower Gp mRNA levels. While at 8hpf and 24hpf the wt PGCs indeed tend to have significantly more Gp mRNAs than the Mmut PGCs, at 3.5hpf no significant difference was found (Figure 2F). In line with this observation, bulk RNA-seq at the 1-cell stage did not reveal consistently lower Gp mRNA levels in Mmut embryos (Figure S2E). Furthermore, transcriptome-wide we find only a very low number of significantly differentially expressed genes at 3.5hpf (Figure S2F). Hence, the reduction in PGC number observed upon loss of maternal Tdrd6a most likely is not due to an overall reduction of Gp mRNA levels or other general transcriptome differences.

We then computed all pairwise correlations between the individual Gp mRNAs in wt PGCs at 3.5hpf. For comparison, we calculated the pairwise correlation of non-Gp background (BG) mRNAs (see experimental procedures). This revealed a general positive correlation for Gp mRNAs in wt PGCs (Figure 2G and Figure S2G), indicating that relatively fixed ratios of individual Gp transcripts are loaded into PGCs. Strikingly, in Mmut PGCs this positive correlation is completely lost (Figure 2H and Figure S2G). Together, these data show that the stoichiometry of Gp mRNAs in single PGCs is tightly controlled and that this depends on maternally provided Tdrd6a.

### Tdrd6a interacts with known Gp mRNAs

Given these results, we next explored whether Tdrd6a interacts with Gp-residing mRNAs via Tdrd6a RNA-IP followed by sequencing (RIP-seq) experiments. Strikingly, all known Gp mRNAs (Hashimoto et al., 2004; Koprunner et al., 2001; Strasser et al., 2008; Wang et al., 2013; Weidinger et al., 2003; Yoon et al., 1997) were strongly enriched in the Tdrd6a RIP-seq compared to input (Figure 3A). We validated these findings using Tdrd6a RIP-qPCR for the Gp markers *vasa*, *dazl*, and *nanos3*, revealing between fifty- and a hundred-fold enrichment in the Tdrd6a RIPs (Figure 3B). The mRNA that was most strongly enriched in the RIP-seq was *hook2* (Figure 3A), coding for a microtubule binding protein. *Hook2* is an unknown Gp component in zebrafish, but reported to be present in *Xenopus* Gp (Owens et al., 2017). Interestingly, in our scRNA-seq data, *hook2* behaves similar to other Gp markers and also displays the typical Tdrd6a-dependent positive correlation with other Gp transcripts (Figure S3A and S3B). Indeed, *in situ* hybridization (ISH) confirmed presence of *hook2* in zebrafish Gp (Figure 3C). Finally, translation inhibition morpholino (MO) injections to knock down *hook2* revealed the requirement of *hook2* for the generation of normal PGC numbers (Figure 3D) substantiating our findings further that *hook2* is a *bona fide* Gp component, relevant for germ cell formation.

**Figure 3.**
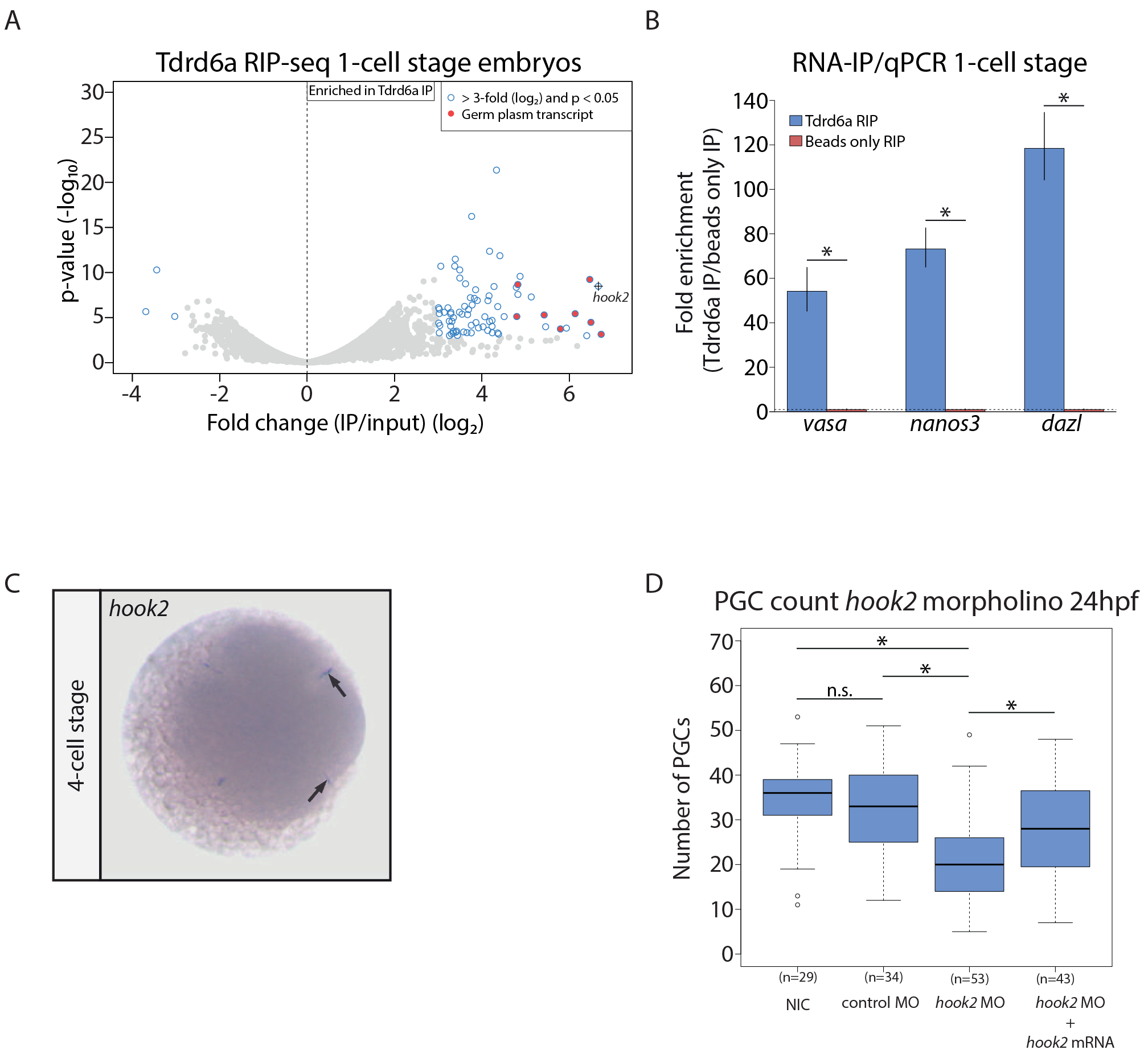
Tdrd6a interacts with mRNA. (A) Vulcanoplot displaying the fold difference between Tdrd6a RIP-seq and input on the x-axis (average of three biological replicates). Y-axis indicates the p-value belonging to the observed differences between Tdrd6a RIP-seq and input. (B) Tdrd6a RIP-qPCR analysis for *nanos3*, *dazl* and *vasa*. Enrichments were calculated compared to beads only RIP and normalized with *β-actin*. Error bars represent standard deviation of two biological replicates (* indicates p-value < 0.001, p-value obtained by two-sided Student’s t-test). (C) ISH against *hook2* at the 4-cell stage. Arrows indicate Gp. (D) Quantification of PGC numbers observed in embryos in morpholino knock-down (MO KD) injection experiment. NIC = non-injected control, the control MO targets the *fus* transcript, the *hook2* mRNA contained mismatches at the *hook2* MO target site and could rescue the KD (* indicates p-value < 0.01, n.s. indicates non-significant, calculated by Wilcoxon test).

It has been proposed that in *Drosophila*, the PIWI protein Aub binds Gp mRNAs in embryos and regulates their stability and localization to Gp (Barckmann et al., 2015; Vourekas et al., 2016). In analogy, we performed Ziwi RIP-qPCR experiments on 1-cell stage embryos for *vasa*, *dazl* and *nanos3*, using the same experimental conditions used for the Tdrd6a RIP-qPCR experiment. The enrichment values for the tested mRNAs were all below three-fold (Figure S3C), while Western blot confirmed that the IPs were successful (Figure S3D). These enrichment values in the Ziwi-RIPs are in sharp contrast to the values obtained in Tdrd6a RIPs. We conclude that in zebrafish, Tdrd6a strongly associates with Gp mRNAs in a Ziwi-independent manner.

### Tdrd6a is essential for Bb and Gp integrity

We next asked whether Tdrd6a affects the localization of Gp-residing mRNAs. First, we performed whole mount fluorescent ISH (FISH) on oocytes against *dazl*. *Tdrd6a* mutant oocytes often display aberrant staining patterns. For instance, the Bb often appeared to be smaller relative to the entire oocyte and lacked a well-defined edge or was even further distorted (Figure S4A, left panels). We quantified these defects by classifying the observed structural abnormalities (Figure S4A) and calculating the size ratio between the Bb and the oocyte, which is significantly smaller without Tdrd6a, irrespective of the probe used (Figure S4B).

We extended these experiments by combining double single-molecule FISH (smFISH) with IHC for Tdrd6a in a Buc-eGFP positive background (Riemer et al., 2015). In the Bb, Buc-eGFP forms a continuous structure, establishing an ordered matrix in which other components can be embedded (Figure 4A). Tdrd6a overlaps with the Buc fibers, even though it is more punctate. The smFISH signals show that different Gp transcripts display diverse sub-localization to the Bb. The *dazl* signal is found as a rather compact core of the Bb, whereas *vasa* is found more throughout the entire Bb (Figure 4A). Interestingly, the smFISH signals seem not to overlap with each other, but rather form a transcript-specific network (Figure 4A, line graph).

**Figure 4.**
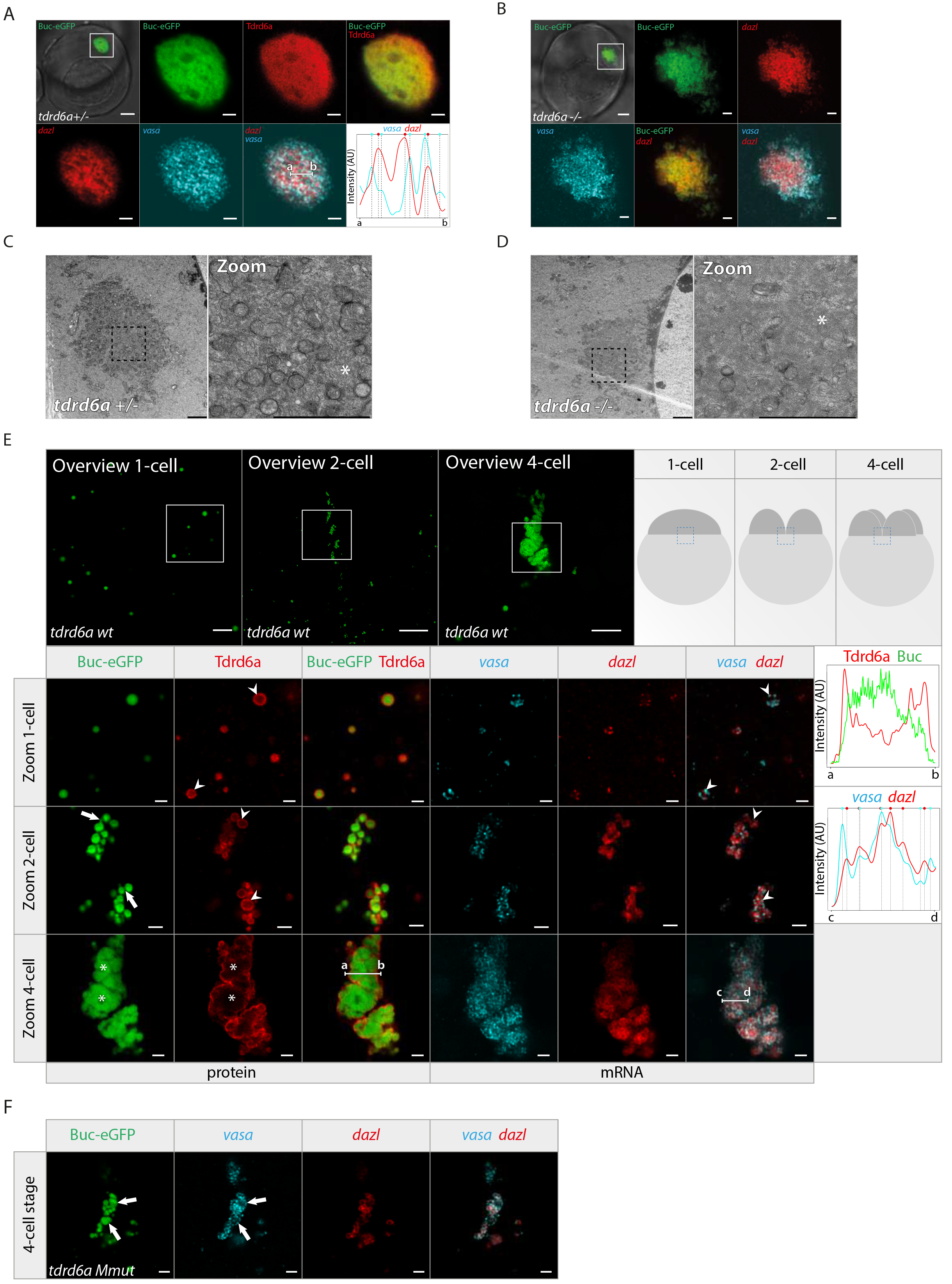
The integrity of the Bb and Gp depends on Tdrd6a. (A, B) Confocal images of Buc-eGFP positive oocytes in *tdrd6a +/−* (A) and *tdrd6a −/−* (B) background. IHC for Tdrd6a and double smFISH was performed and displayed as indicated. Tdrd6a and Buc overlap, whereas *dazl* and *vasa* typically do not. Line graph displays intensity for *dazl* (red) and *vasa* (cyan) signals over line a-b (see overlay), with vertical lines indicating fluorescence peaks per smFISH signal (highlighted by colored circle on top) showing transcript peaks are in a separate phase. (C,D) Electron micrographs of *tdrd6a +/−* (C) and *tdrd6a −/−* (D) oocytes displaying the Bb (electron-dense area). Asterisks indicate more homogeneity in the mutant Bb compared to the heterozygous situation. (E) Confocal images of Buc-eGFP positive embryos from *tdrd6a +/−* mothers, at 1-,2- and 4-cell stage overview, focusing on Gp as schematically indicated. In zoom of the Gp, IHC for Tdrd6a and double smFISH was performed and displayed as indicated. Arrowheads indicate mRNA peripherally localized on the Gp-droplet, arrows indicate Buc-eGFP bridges, asterisks mark mature Gp, containing fused Buc-eGFP and mRNA accumulated inside of the structure. Line graphs display intensity for Buc (green) versus Tdrd6a (red) signals over line a-b and *dazl* (red) and *vasa* (cyan) intensity over line c-d, with vertical lines indicating fluorescence peaks per smFISH signal (highlighted by colored circle on top). (F) Confocal images of Buc-eGFP positive Gp of a 4-cell stage embryo of a *tdrd6a −/−* mother (Mmut). SmFISH was performed and displayed as indicated. Arrows indicate areas where Buc-eGFP has fused, but mRNA remains peripherally localized. Scalebars indicate 10μm (overview), 2μm (zoom) (A,B,E,F) and 2μm (C,D).

In *tdrd6a* mutant oocytes, the Bb marked by Buc-eGFP is more irregular and lacks a constricted outline, confirming our previous observations (Figure 4B). Gp-transcripts still localize to the Bb but due to the scattered nature, organization as found in heterozygous oocytes is lost (Figure 4B).

A more widely conserved function of the Bb is mitochondrial selection, which are therefore highly represented in the Bb (Bilinski et al., 2017). In *Xenopus*, they are embedded in a matrix formed by Xvelo (Boke et al., 2016). We show this is also the case in zebrafish, using a mitotracker in the Buc-eGFP background (Figure S4C). In *tdrd6a* mutant oocytes, mitochondria still localize to the Bb, though their organized embedding seems less distinct (Figure S4C). Electron micrographs could confirm that mitochondria are still recruited to the Bb in the absence of Tdrd6a, however, Bb organization loses heterogeneity, and electron-dense areas are more monotonous (Figure 4C/D and S4D, asterisks). In conclusion, Tdrd6a is required for the overall organization of the Bb, possibly through incorporation and/or stabilization of Bb components.

### Tdrd6a is required for merging particles with distinct mRNA content into mature Gp structures

Upon fertilization, rapid changes occur in the Gp during the first divisions. Small assemblies containing Buc accumulate and fuse at the cleavage planes into larger Gp structures (Riemer et al., 2015). We therefore performed smFISH and IHC experiments on 1-cell, 2-cell and 4-cell stage embryos, to capture the different steps during this reorganization. This revealed that in 1-cell stage embryos, Buc and Tdrd6a form isolated particles, decorated with discrete mRNA foci at their periphery (Figure 4E). Buc forms the core of the Gp particles, whereas the Tdrd6a signal surrounds them. Interestingly, we again observe that transcript signals do not overlap, even though they do co-exist on Gp droplets. Amounts per transcript can vary per aggregate (Figure 4E), possibly due to stochastic variation. At the 2-cell stage, the droplets organize themselves along the cleavage plane and start to cluster together (Figure 4E). The Buc signal often bridges individual granules, but they do not fuse (Figure 4E, arrows). Furthermore, Tdrd6a appears to be localized around the Buc-assemblies, similar to the mRNA (Figure 4E, arrowheads). The Gp further grows towards the 4-cell stage into a larger structure, in which the Tdrd6a surrounds the collected Gp (Figure 4E, asterisks). We also observe that in these parts, the mRNA has mostly moved inwards, forming large, intermingled networks (Figure 4E, line graph), which co-occurs with enlarged Buc aggregates (Figure 4E, asterisks). Overall, the smFISH signals for different mRNAs are very well mixed within the larger Gp structure, but areas of overall enrichment for one or the other mRNA can still be observed.

The various Gp structures do not display an internal Tdrd6a signal. FISH studies using DIG-labelled probes followed by IHC against DIG show that primary and secondary antibodies are clearly capable of entering the inner part of the Gp (Figure S4E), suggesting Tdrd6a is truly enriched at the periphery. We note, however, that the apparent peripheral localization of Tdrd6a based on IHC could also be due to very high Tdrd6a concentrations on the outside of the Gp, preventing sufficient penetration of specifically the Tdrd6a antibody into the structure.

In *tdrd6a* MZ embryos, mRNAs still localize to the Gp particles (Figure 4F). However, the structure fails to mature and remains relatively small and highly fragmented, with mRNAs surrounding Buc (Figure 4F and S4F). We do observe some apparent fusion of Buc particles, but typically, also in these cases mRNA remains at the periphery (Figure 4F, arrows). These observations lead us to propose that Gp forms through the ongoing accumulation of small granules, containing Buc, Tdrd6a and mRNPs. Upon maturation, Buc granules merge and the mRNP particles are moving inwards and start to intermingle, forming networks with mRNPs of the same kind. The formation of these larger aggregates depends on Tdrd6a, since mRNPs can still associate with Buc in its absence, but fail to accumulate into mature Gp.

### Tdrd6a directly interacts with Buc via symmetrically dimethylated arginines

The IP-MS experiments on ovary extracts identified Buc as a strong interactor of Tdrd6a (Figure 1D). We also found Buc to be among the strongest interactors of Tdrd6a in freshly laid embryos (Figure 5A). We verified this interaction on Western blot and showed it is resistant to RNase A treatment (Figure S5B). Protein-protein interactions with Tdrds often depend on symmetrically dimethylated arginine (sDMA) residues in its binding partner (Siomi et al., 2010). Indeed, analysis of our MS results identified two dimethylated arginine residues within the C-terminus of Buc, residing in a tri-RG (RG(X0-4)RG(X0-4)RG) motif (Figure 5B) (Thandapani et al., 2013). In order to test their relevance for interaction with Tdrd6a, we performed pull-down experiments using biotinylated peptides covering these arginines in either an sDMA-or non-methylated state (Figure S5B) followed by MS. In the pull-down using the methylated Buc-peptide, Tdrd6a was highly enriched (Figure 5C). The pull-down with the non-methylated peptide showed enrichment for two members of the serine/arginine-rich protein kinase complex Srpk1a and Srpk1b (Figure 5C), confirming the pull-down was successful and revealing potential additional post-translational regulation of Buc besides sDMAs. Moreover, further pull-down experiments demonstrated that all three sDMAs are required for Tdrd6a interaction, since single or double sDMAs did not show this enrichment (Figure S5C).

**Figure 5.**
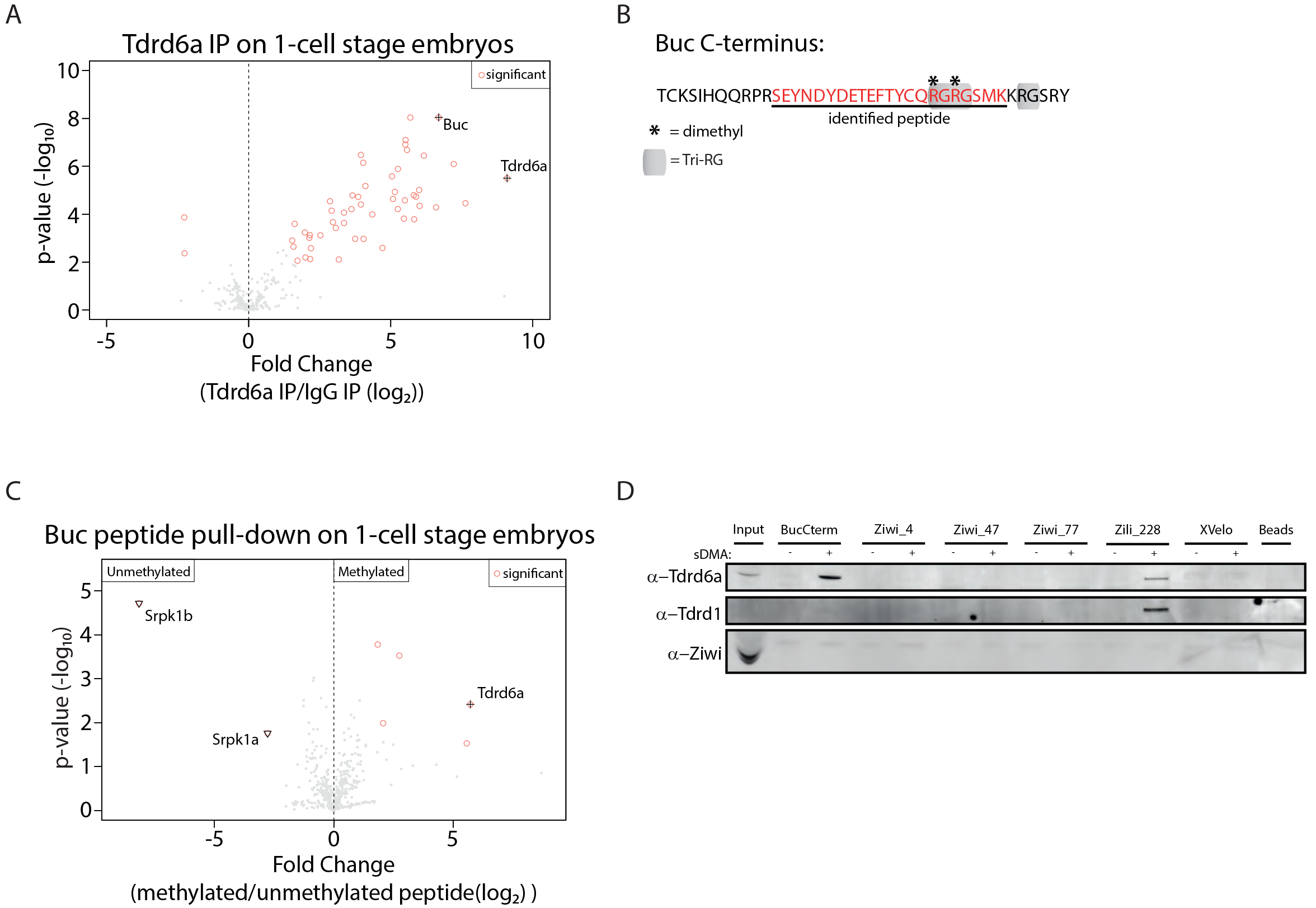
Tdrd6a and Buc interact via sDMAs in the C-terminus of Buc. (A) Volcanoplot of Tdrd6a IP compared to IgG IP on embryo extracts, followed by MS. (B) C-terminus of Buc with the identified dimethylated peptide underlined in red. 3 RG sites together form a tri-RG motif, indicated in grey. (C) Volcanoplot of peptide pull-down on embryo extracts followed by MS. Peptides were synthesized based on Buc C-terminal region. The ‘Methylated’ peptide contained 3 sDMAs and was compared to an unmodified version of the same sequence (‘Unmethylated’) (Also see Figure S5B). (D) Peptide pull-down followed by Western for multiple methylated (sDMA) and unmethylated peptides derived from proteins known to contain sDMA modifications and the Buc homolog XVelo on ovary extracts.

To test the specificity of the Buc peptide for Tdrd6a we repeated the pull-down, taking along additional peptides derived from Ziwi, Zili and XVelo, either sDMA-modified or non-methylated. On Western blot, these experiments showed clear enrichment of Tdrd6a using the methylated Buc peptide, but not using the non-methylated Buc peptide (Figure 5D). We do see some affinity of Tdrd6a for the methylated Zili peptide previously shown to interact with Tdrd1 (Huang et al., 2011). Since Zili is not maternally provided, this affinity could be biologically relevant in ovarian nuage where Zili interacts with Tdrd6a (Figure S1G). In turn, Tdrd1 displayed affinity only for the methylated Zili228 peptide, as expected, and did not interact with the sDMAs residing in the Buc C-terminus (Figure 5D). We conclude that Tdrd6a specifically interacts with the sDMA-modified C-terminal part of Buc, and demonstrate potential sequence specificity in sDMA-Tudor domain interactions in general.

### Tdrd6a affects the aggregation behavior of Buc

It has been shown previously that XVelo, the homolog of Buc, is a protein with a prionlike domain at its N-terminus that has a tendency to self-aggregate. We showed that Buc aggregation, illustrated by detailed Bb and Gp imaging, is highly regulated *in vivo*. Consequently, we reasoned that Tdrd6a might be involved in spatio-temporal regulation of Buc aggregation. We first tested this in a heterologous cell culture system, BmN4 cells. These are ovary-derived and kept at 27°C, thereby mimicking natural conditions for Tdrd6a and Buc. Expression of Buc results in abundant, cytoplasmic, small granules (Figure 6A). In contrast, Tdrd6a displays a ubiquitous cytoplasmic signal (Figure 6B). Co-transfection of both proteins results in dramatic changes, with two possible outcomes: Either Tdrd6a and Buc are found to co-localize in enlarged, cytoplasmic aggregates, or both Tdrd6a and Buc display diffuse cytoplasmic localization (Figure 6C and D, respectively). We then repeated the experiment, but performed consecutive-, rather than co-transfection and quantified protein behavior. If we first transfect Buc, followed by Tdrd6a on the next day, we always observed enlarged granules that are positive for both Buc and Tdrd6a (quantification shown in Figure 6E). When Tdrd6a is transfected first, Buc is mostly localizing throughout the cytoplasm (Figure 6F). Only when the Tdrd6a signal is low, Buc seems to be able to form enlarged granules. This effect of Tdrd6a on Buc is specific to Tdrd6a, since co-transfection with for example Dcp1, a P-body marker, leaves both proteins unaffected (Figure S6A, S6B). These results demonstrate that Tdrd6a can either stimulate the accumulation of Buc into larger granules or prevent its aggregation altogether.

**Figure 6.**
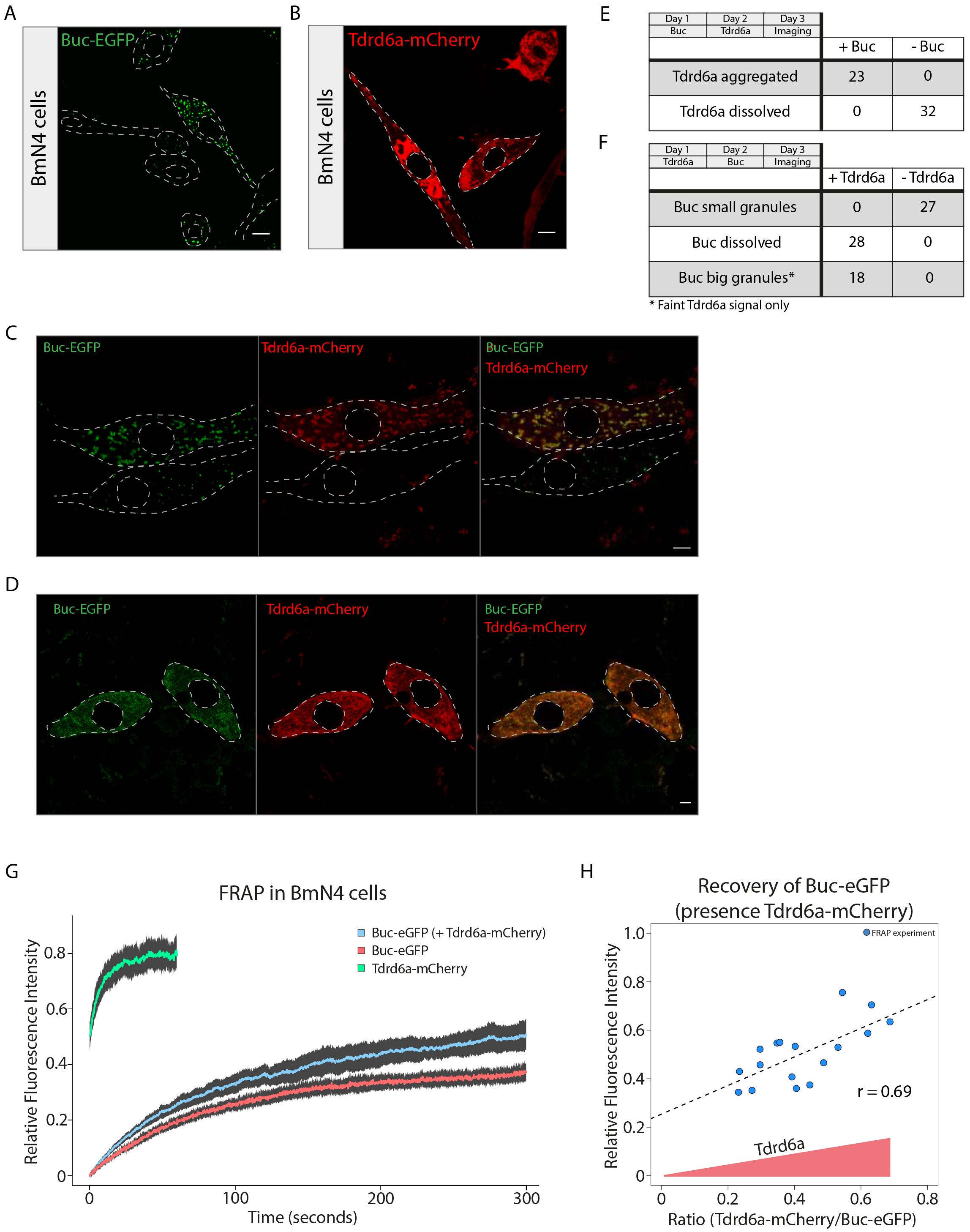
Tdrd6a stimulates Buc mobility in BmN4 cells. (A) Localization of Buc-eGFP 1dpt. (B) Localization of Tdrd6a-mCherry in BmN4 cells 1dpt. (C, D) Cotransfected BmN4 cells displaying enlarged granules in Tdrd6a-mCherry positive cell (upper cell, C) but not in negative cell (lower cell, C), or dissolved Buc-eGFP (D). (E) Quantification of localization of Tdrd6a transfected 1 day after Buc. (F) Quantification of localization of Buc, transfected 1 day after Tdrd6a. (G) FRAP recovery curves of Tdrd6a-mCherry and Buc-eGFP (with or without the presence of Tdrd6a-mCherry as indicated). Fluorescence intensity is the calculated fraction of the pre-bleach intensity, and plotted with the 95% confidence interval. Tdrd6a-mCherry reaches a plateau quickly upon bleaching (60 seconds recovery plotted), compared to Buc-eGFP (300 seconds recovery). (H) FRAP recovery of Buc-eGFP plotted against increasing relative amounts of Tdrd6a-mCherry present in the bleached granule. Scale bars: 10μm (A,B), 5μm (C,D).

We then continued to investigate the physiological properties of these granules in more detail using Fluorescence Recovery After Photobleaching (FRAP). Tdrd6a recovers rapidly upon bleaching, reflecting a high mobility in and out of the granule (Figure 6G, n=17). Interestingly, while Buc alone recovers up to ~35% (n=17) of the initial fluorescence intensity, this drastically increases in the presence of Tdrd6a to ~55% (n=17) (Figure 6G). Quantification of the FRAP experiments shows that the increase in Buc recovery in the presence of Tdrd6a is significant (Figure S6C). However, we did observe a rather broad distribution in recovery in Tdrd6a-Buc double positive granules (Figure S6C). We hypothesized that this variation in recovery could be due to differences in relative protein concentrations in the individual granules that are studied. Hence, we normalized the protein amounts in the FRAP experiments by calibrating relative fluorescence using an mCherry-eGFP construct (Figure S6D). This revealed that the more Tdrd6a is present in the granule, the better Buc can recover (Figure 6H). Furthermore, in lysates of transfected BmN4 cells, Buc-eGFP cannot be detected in the soluble fraction in the absence of Tdrd6a-mCherry (Figure S6E and S6F). In agreement with our FRAP results, we found significant amounts of Buc-eGFP in the soluble fraction of BmN4 lysates when Tdrd6a-mCherry is present as well (Figure S6E). We conclude that Tdrd6a positively stimulates Buc mobility and solubility.

Lastly, to investigate the *in vivo* relevance of these results, we performed FRAP on the Bb of Buc-eGFP positive oocytes in the *tdrd6a +/−* and *−/−* background. Indeed, these FRAP studies also show the remarkable decrease of mobility of the Bb in the absence of Tdrd6a (Figure 7A). In conclusion, our data suggest that Tdrd6a is a potent affector of the aggregating behavior of Buc-containing structures and stimulates their growth, heterogeneity and mobility (Figure 7B). We could demonstrate this both *ex vivo* in transfected BmN4 cells as well as *in vivo* in a complex and poorly understood, phase-separated structure like the Balbiani body.

**Figure 7.**
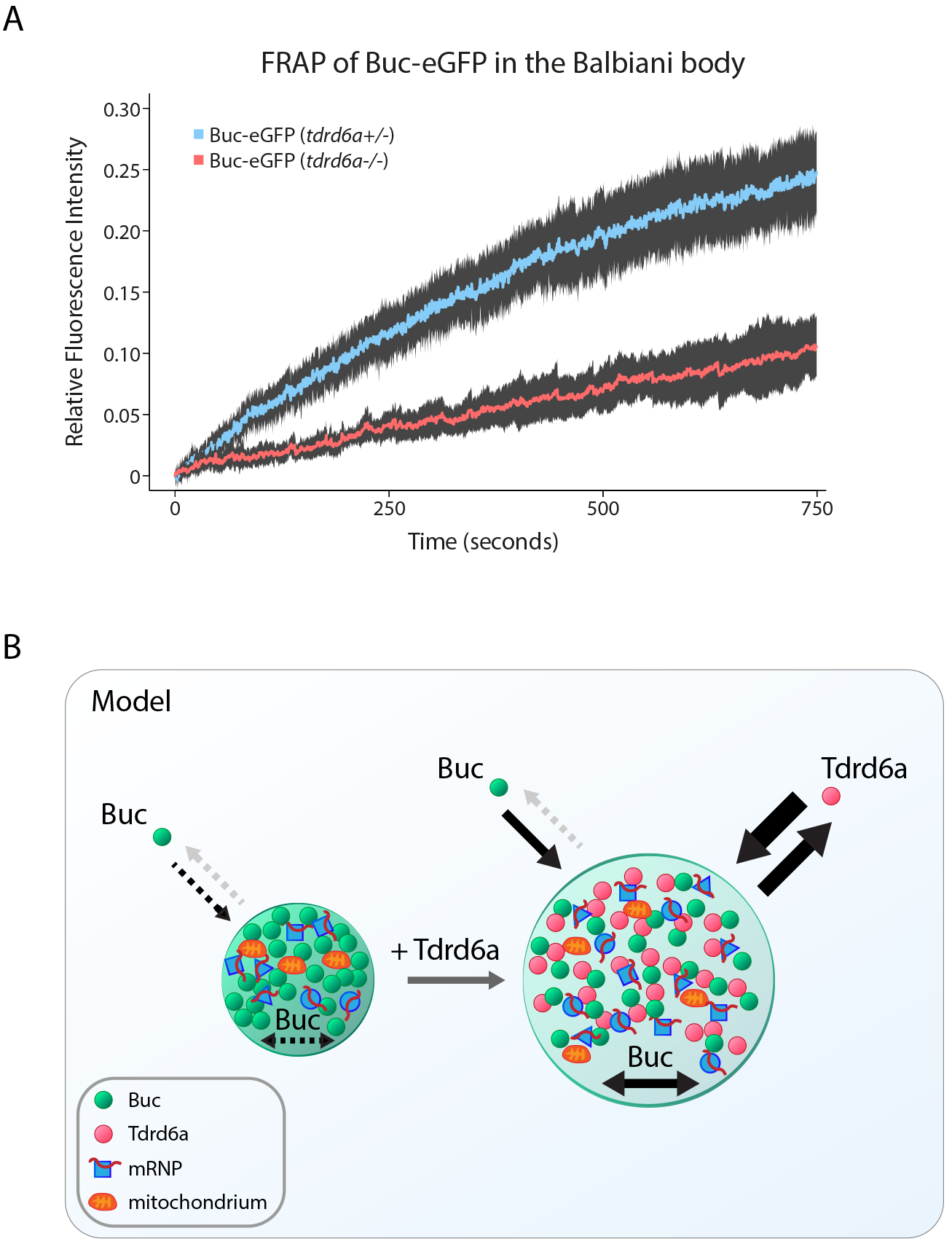
Tdrd6a stimulates Buc-eGFP mobility *in vivo*. (A) FRAP recovery curves of Buc-eGFP in *tdrd6a* heterozygous or mutant Balbiani bodies. Fluorescence intensity is the calculated fraction of the pre-bleach intensity and plotted with the 95% confidence interval. (B) Model of Buc-containing granules, with or without Tdrd6a. Arrows indicate movement in and out of the droplets or mobility within the droplet itself.

## Discussion

Proteins like Tdrd6a, with multiple Tudor domains in tandem, are well known to act in germ cells, in particular in small RNA pathways and their organization in perinuclear aggregates or granules. Their precise molecular functions, however, are far from resolved. Another type of protein that is often found in germ cells has low-complexity regions and/or prion-like domains, such as Buc. Other examples are MUT-16 and MEG proteins from *C. elegans* and Xvelo from *Xenopus*, which nucleate a variety of subcellular aggregates (Boke et al., 2016; Phillips et al., 2012; Wang et al., 2014). However, insights into how their aggregation behavior is regulated remain scarce. We demonstrate that Tdrd6a regulates the aggregation Gp-related structures. More specifically, it promotes solubility and mobility of Buc, and thereby the assembly of Buc-driven aggregates into larger structures containing well-determined amounts of germ cell-specifying mRNAs and other Gp components. Various aspects related to our findings will be further discussed here.

### Tdrd6a does not affect piRNA generation

Even though the Piwi protein Ziwi is one of the strong interactors of Tdrd6a in ovary extracts, lack of Tdrd6a does not have an effect on piRNA accumulation. Given the intimate connection between piRNA biogenesis and function, a mechanistic role for Tdrd6a in the piRNA pathway does not seem likely. Only during early oogenesis, we observe a loss of antisense bias in the piRNA population. Since Ziwi predominantly interacts with antisense piRNAs (Houwing et al., 2007), this observation may reflect a transient role for Tdrd6a in Ziwi accumulation and/or stabilization during early oogenesis.

### Molecular basis behind the PGC phenotype of *tdrd6a* mutants

We observed that in embryos, Gp arises from the continuous merging of smaller Buc-Tdrd6a units. In these granules, Buc is always found at the core, while we detected Tdrd6a mainly at the periphery. Possibly, however, Tdrd6a is also present within the core. The individual Buc-Tdrd6a granules in 1-2 cell embryos display discrete mRNA foci at their circumference, as visualized by smFISH. Interestingly, these foci move more internally when larger assemblies arise, where they start to form networks. Homotypic assemblies of mRNPs have been recently described in *Drosophila*, where it has been demonstrated that Gp mRNAs initially form homogenous mRNP granules, followed by fusion into heterogeneous mRNP aggregates, in which the quantities of Gp mRNAs positively correlate (Little et al., 2015; Trcek et al., 2015). In zebrafish, we could infer that mRNA quantities in mature Gp positively correlate as well, based on PGC scRNA-seq. In order for this correlation to be established, Tdrd6a is critical. We note that in our experiments we select for successfully formed PGCs through FACS, and are blind to PGCs that may have failed to become specified. Therefore, the effects we report in our scRNA-seq experiments most likely underestimate the impact of Tdrd6a on PGC development.

Why does this correlation between Gp mRNAs depend on Tdrd6a? In absence of Tdrd6a, even though transcript levels deposited by the mother remain stable, mostly small, incomplete Gp-like structures are found. Given that each Tdrd6a-Buc granule at the 1-2 cell stage only has a limited number of individual mRNPs, a sufficiently large number of Buc-Tdrd6a granules needs to accumulate to reach the stable ratios in the final Gp as would be found overall on all sub-granules combined. It has been suggested previously that Buc-containing particles behave like liquid droplets (Riemer et al., 2015). Without Tdrd6a, the number of granules that merge is low, potentially resulting in skewed relative abundances of mRNPs in the relatively small Buc assemblies, simply because of stochastic fluctuation. We propose that Tdrd6a affects Buc aggregation behavior such, that the smaller individual Buc aggregates become less rigid and more liquid-like, thereby enhancing fusion into larger, complete Gp assemblies. Since single Gp components can have a strong impact on the efficiency of PGC formation, something we again observe with the newly identified zebrafish Gp mRNA component *hook2*, this reduction in Buc aggregation may directly relate to the observed PGC specification and/or maintenance defects.

### Potential intrinsic characteristics of mRNPs dictate subgranular organization

Single Buc-containing particles contain individual foci of various Gp transcripts at their periphery. In more enlarged Gp structures, however, we observe bigger networks consisting of one type of mRNA that spread throughout the Gp and do not remain at the periphery. We speculate that this transition may take place as follows. Initially, mRNPs are displayed at the periphery of the Gp droplets marked by Buc. This is Tdrd6a independent, since recruitment also occurs in *tdrd6a* mutants. Tdrd6a is highly concentrated around these granules and as we demonstrate, increases Buc mobility. This effect of Tdrd6a on Buc may lead to the formation of the bridges of Buc protein between individual Buc granules. These bridges, and later on the more extensive fusion of Buc granules, as demonstrated by Riemer *et al.* (Riemer et al., 2015), brings together a much larger number of individual mRNPs. When transcripts accumulate to high enough concentrations, their intrinsic properties may trigger the formation of larger homotypic clusters and/or networks within the Gp structure. Indeed, intrinsic tendency of transcripts of the same kind to cluster is a phenomenon that has been suggested previously for *Drosophila* Gp (Little et al., 2015; Trcek et al., 2015). These larger homotypic structures may be stabilized further by their continued interaction with the growing Buc-network, in which they intermingle with other homotypic networks. Whether the interactions between Buc and these mRNPs is direct or not is currently unclear, but it has been speculated that the C-terminal region of Xvelo, a region analogous to the interaction site of Buc with Tdrd6a, can interact with RNA (Boke et al., 2016). Whether indeed Tdrd6a and mRNPs compete for overlapping binding sites on Buc remains to be determined. Finally, it is currently still unclear why certain mRNPs localize to Gp in the first place. Intrinsic sequences of the mRNAs may play a role (Knaut et al., 2002; Koprunner et al., 2001), but the fact that we identify the cytoplasmic EJC complex and the CPEB complex in our Tdrd6a interactome may reveal an additional aspect: mRNPs that have not undergone any translation could be prone to be incorporated into Gp-related structures. Interestingly, these complexes have been demonstrated in *Drosophila* and *Xenopus* to play a role in translational control and/or Gp transcript localization (Hachet and Ephrussi, 2004; Minshall et al., 2007; Nelson et al., 2004). Possibly this, in combination with inherent properties to accumulate into larger homotypic assemblies is key to understand the specificity of mRNP incorporation into Gp structures.

### Buc-Tdrd6a interaction as a model for regulated protein aggregation

The assembly of proteins with intrinsically disordered regions, such as Buc, has been recognized as a research field of major importance. First, it represents a novel type of compartmentalization, which may steer biochemical processes. There appears to be a rather wide range of aggregation states, spanning from liquid-like droplets to almost solid aggregations (Brangwynne et al., 2011; Patel et al., 2015; Shin and Brangwynne, 2017). It has been proposed that pathogenic protein aggregates, such as found in Alzheimer’s disease or ALS, represent an exaggerated state of a normally occurring, aggregating form of a protein. Hence, knowledge about how aggregation states can be regulated will be directly relevant to the understanding of these types of disease. Buc aggregation is very dynamic during zebrafish oogenesis and embryogenesis. It starts by forming a very stable Bb, then disperses into fragments and upon fertilization forms Gp along the first cleavage planes. Finally, within early PGCs, Buc can be found in peri-nuclear granules. We demonstrate that the aggregation behavior of Buc is strongly affected by another protein, Tdrd6a, and that this regulatory potential may be affected by PTMs, in this case arginine di-methylation. During early embryogenesis in *C. elegans*, phosphorylation and dephosphorylation of MEG-1/3 control P-granule disassembly and assembly, respectively (Wang et al., 2014). Interestingly, we find specific kinases to be enriched in the non-modified Buc peptide pulldown making it tempting to speculate that besides arginine dimethylation, also phosphorylation may regulate Buc aggregation properties. The Buc-Tdrd6a system that we characterize both *in vivo* and *ex vivo* represents a powerful model to study the spatio-temporal regulation of protein aggregation, both by trans-acting factors as well as PTMs.

Finally, our work suggests that Tdrd6a, and potentially other multi-tudor domain proteins do not themselves act as scaffolds for aggregation. Rather, Tdrd6a affects the aggregation behavior of another protein that in turn acts as a scaffold for aggregation. Analogous to previous studies, Buc typically behaves like a ‘scaffold’: recovering slowly and only partially. Tdrd6a recovery is typical for a granule ‘client’, displaying rapid, near complete recovery, indicating high mobility in and out of the Buc-aggregate (Woodruff et al., 2017). This may mean that in other scenarios in which Tdrd6a-like proteins have been described to affect aggregations, such as for example the chromatoid body in mammalian spermatocytes or peri-nuclear nuage, scaffold proteins such as Buc are still to be discovered. Alternatively, well-known proteins may in fact act as such scaffolds, without us currently realizing this. For instance, Piwi proteins typically have rather long and seemingly unstructured N-terminal tails, and also other well-studied proteins like for example Vasa contain disordered regions. It is therefore possible that such proteins act as aggregation scaffolds during assembly of peri-nuclear nuage.

## Experimental procedures

### Whole mount smFISH and IHC

PFA fixed oocytes/embryos were collected as described above and were incubated overnight with 1:100 anti-Tdrd6a and 1:100 of both smFISH probes (Stellaris™ custom design) stocks (12.5μM in TE buffer) in hybridization buffer (10% dextran sulfate, 10% formamide, 1mg/mL tRNA, 0.02%BSA, 2mM vanadyl-ribonucleoside complex (NEB S1402S) in 2xSSC) at 30oC. Next day, wash 15 minutes in wash buffer (10% formamide, 2xSSC) and incubate in 1:500 anti-rabbit alexa-405 for 30 minutes in wash buffer. Then wash 2 × 15 minutes in wash buffer and mount in ProLong™ Gold Antifade Mountant (ThermoFisher).

### Peptide pulldown

Peptides were synthesized by Peptide Specialty Laboratories GmbH (for peptide sequences see supplemental experimental procedures). 20μg peptide in 500μL IP buffer (25mM Tris pH 7.5, 150mM NaCl, 1.5mM MgCL2, 1% Triton-X100, 1mM DTT) was pre-incubated with streptavidin-coupled magnetic beads for 30 minutes at RT, rotating. 20μL resin (ThermoFisher 65001) was used per pulldown. Then, the respective lysate was added to the washed beads and incubated for 1hr at 4oC while rotating. The beads were then washed with wash buffer (25mM Tris pH 7.5, 300mM NaCl, 1.5mM MgCL2, 1mM DTT) and either used for Western blot analysis or MS.

### FRAP

FRAP was performed on a TCS SP5 Leica confocal microscope, equipped with a FRAP-booster, using a 63× oil objective with an NA of 1.4 (BmN4 cells) or a 63× water objective with an NA of 1.2 (Bb). In BmN4 cells, entire granules were bleached in a fixed region of 1.5μm ø and recovery was followed for 1500 frames (0.2s/frame). Bbs were bleached partially in a fixed region of 2.5μm ø and recovery was followed for 1500 frames (0.5s/frame). Regions (pre- and post-bleach) were tracked using TrackMate (Tinevez et al., 2017). 10 pre-bleach frames were recorded and after background subtraction, the average intensity was used as pre-bleach intensity. Postbleach frames were background subtracted and to make replicates comparable, postbleach frame #1 of each measurement was set to 0 and corresponding pre-bleach intensity was corrected for this. Normalization of Buc-eGFP and Tdrd6a-mCherry was performed by plotting intensities of mCherry-eGFP using the same microscope settings as for the FRAP. Intensities were plotted after background subtraction and the resulting curve was used to calculate protein ratios using the initial/pre-bleach intensities of each experiment.

## Accession numbers

The accession number for the raw and processed RNAseq data is GSE79285. The mass spectrometry proteomics data have been deposited to the ProteomeXchange Consortium via PRIDE partner repository with the dataset identifier PXD008322.

## Supplemental Information

Supplemental information consists of additional Experimental Procedures and six supplemental figures.

## Author contributions

LJTK, EFR and RFK conceived the study and designed experiments. LJTK performed scRNA-seq experiments and data analysis, RIP-seq analysis and IP experiments. AMJD performed smRNA-seq analysis. EFR performed most other experiments. HH performed initial analyses of the *tdrd6a* mutant and Tdrd6a localization. AWB performed BmN4 cell culture experiments. KW performed scRNA-seq experiments and DG devised scRNA-seq data analyses. SR performed electron microscopy experiments and CTH optimized RIP-seq protocols. FB performed MS analysis. AvO and RFK supervised the project. LJTK, EFR and RFK wrote the paper with input from all authors.

## Acknowledgements

We thank the members of our laboratory for fruitful discussions. Yasmin El Sherif is thanked for extensive experimental support. We thank the Cuppen group (Hubrecht) for identifying the *tdrd6a*^*Q158X*^ allele. We would like to thank Jeroen Krijsgveld and the proteomics core facility at EMBL for performing initial proteomics analysis and Eugene Berezikov (Hubrecht/ERIBA) for initial bioinformatics support. We thank the following IMB Core Facilities for their contributions and valuable services: Genomics, Microscopy, Bioinformatics, Flow Cytometry, and the Media Lab. In particular, we thank Mária Hanulová for support in FRAP experiments. We thank Edward Lemke for critical reading of the manuscript. This work was supported by the Rhineland Palatinate Forschungsschwerpunkt GeneRED, a Marie Curie fellowship 623119 (LJTK), the ERC (ERC-StG202819, RFK, and ERC-AdG294325-GeneNoiseControl, AvO) and Vici awards (RFK, AvO) from the Nederlandse Organisatie voor Wetenschappelijk Onderzoek (NWO).

